# Yeast proteins reversibly aggregate like amphiphilic molecules

**DOI:** 10.1101/2021.03.12.435082

**Authors:** Pouria Dasmeh, Andreas Wagner

## Abstract

More than a hundred proteins in yeast reversibly aggregate and phase-separate in response to various stressors, such as nutrient depletion and heat shock. We know little about the sequence and structural features behind this ability, which has not been characterized on a proteome-wide level. To identify the distinctive features of aggregation-prone regions, we apply machine learning algorithms to genome-scale limited proteolysis-mass spectrometry data from 96 yeast proteins that phase-separate upon heat shock. We find that the aggregation-prone regions (APRs) of our study proteins are significantly enriched in aliphatic residues and depleted in positively charged amino acids. Aggregator proteins with longer APRs show a greater propensity to aggregate, a relationship that can be explained by equilibrium statistical thermodynamics. Altogether, our observations suggest that proteome-wide reversible protein aggregation is mediated by sequence-encoded properties. Aggregating proteins resemble supra-molecular amphiphiles, where APRs are the hydrophobic parts, and non-APRs are the hydrophilic parts.

## Body

Proteins can aggregate reversibly or irreversibly. Irreversible aggregation is often pathological, indicates damaged cellular regulation^1–3^, and is involved in multiple diseases such as Alzheimer’s and Parkinson’s diseases^4,5^. Reversible aggregation, however, can be beneficial and help cells survive stressors. For example, in yeast more than a hundred proteins in multiple subcellular compartments form reversible aggregates in response to nutrient starvation, heat shock, or chemical stress^6^. These aggregated proteins are not misfolded or tagged for degradation^7^. Instead, they help increase cells re-initiate growth during recovery from stress by protecting metabolic enzymes from degradation^6–9^.

Reversible protein aggregation is a special case of a widespread phenomenon called protein phase-separation. In this process, a well-mixed solution of proteins demixes into two phases of high and low densities^10^. Proteins can phase separate with other proteins, but also with RNA or DNA molecules, into biomolecular condensates that regulate transcription^11^, chromatin states^12,13^, and RNA metabolism^14^. We know little about the sequence and structural features behind reversible phase separation, which has not been characterized on a proteome-wide level. In this work, we provide such a view and characterize sequence features that can help predict whether proteins will aggregate reversibly.

We took advantage of recent proteome-wide data from limited proteolysis-mass spectrometry (LiP-MS)^20,21^. This method permits the detection of both pronounced and subtle changes that protein and peptide structures experience in response to stressors such as heat and osmotic shock. Recently, Cappelletti *et. al.* characterized structural changes within 96 proteins that reversibly aggregate upon heat shock in yeast^22^. These proteins are also known as aggregators. They have diverse biological functions, such as telomeric DNA-binding, RNA-helicase activity, or ribosome assembly. Because they are functionally diverse, they constitute an excellent dataset to examine sequence features that facilitate reversible protein aggregation. We studied regions within these proteins whose proteolysis resistance changes significantly upon aggregation ^22^. We refer to these regions as aggregation-prone regions.

What are the characteristic sequence features of APRs? To answer this question, we first performed a principal component analysis on the frequency of 20 amino acids in all 270 APRs of yeast aggregators (Dataset S1). As shown in Figure 1A-B, the first and the second principal components only accounted for ~12% and ~10% of the variation. The PCA biplot for these components shows that no specific combinations of few amino acids preferentially occur in APR sequences (Figure 1B), a further indication that APRs do not show the kind of reduced sequence complexity characteristic of disordered regions. We also looked for linear sequence motifs in APRs using DALEL^29^, an algorithm for the exhaustive identification of degenerate sequence motifs, but found no such enriched motifs.

**Figure 1.**
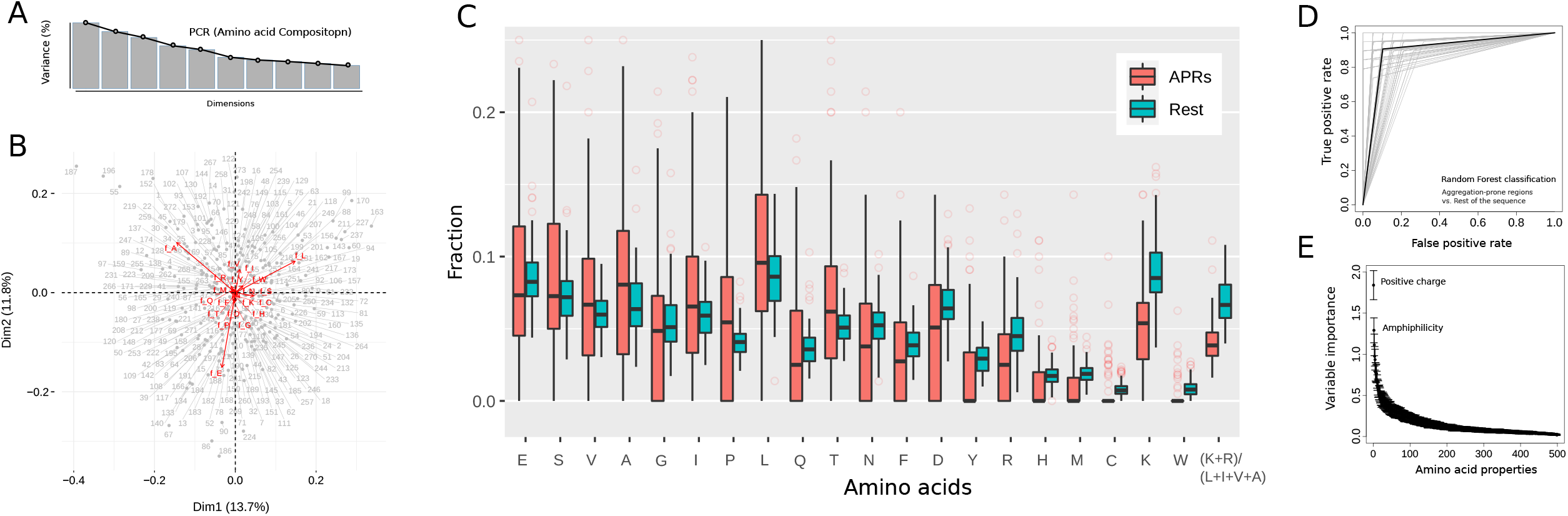
Sequence composition and physicochemical properties of aggregation-prone regions in the yeast aggregators. A) The percentage of explained variance for 10 dimensions from a principal component analysis on the frequencies of 20 amino acids in 270 aggregation-prone regions of 96 yeast proteins. B) Biplot for the first and the second principal components. Grey circles, labeled by numbers from 1 to 270, represent aggregation-prone regions. Red arrows represent the weight of different amino acid frequencies. C) The amino acid frequencies in APRs (shown in red) versus the rest of the protein sequence (shown in green) in 96 yeast aggregators. D) The receiver-operating characteristic (ROC) curves for the clustering of APRs from non-APRs using random forest classification. ROC curves in grey are from 100 random forest clustering and their average is shown in black. E) Ranked importance of physicochemical vartiables quantified as the average decrease in the Gini index for different physicochemical properties.

We then compared the frequencies of amino acids, dipeptides, and tripeptides in APRs with the rest of the protein sequence in aggregators (Dataset S2). Here, we found a significant depletion of the positively charged residues Arg and Lys, and their dipeptides (p~10^−16^; Wilcoxon signed-rank test). We also observed that aliphatic residues Leu, Ile, Ala, Val, were significantly more frequent in APRs compared to the rest of the protein sequence (p~10^−16^; Wilcoxon signed-rank test). The fraction of positively charged to aliphatic residues was the strongest predictor of APRs compared to the fraction of all other amino acids (Figure 1C; p~10^−21^; Wilcoxon signed-rank test).

To find out whether these patterns are linked to differences in physicochemical properties of amino acids, we used a random forest approach, a widely-used machine learning technique for the classification of two or more data sets^30^. Specifically, we used all 96 aggregators with experimentally characterized APRs^22^, and subdivided each protein into two sequence data sets, one comprising the APRs and one comprising the non-APRs. For each of these datasets, we calculated a feature matrix that consisted of 500 amino acid properties (Dataset S3, and S4). We took these properties from the AAindex database^31^, which curates various physicochemical and biochemical properties of amino acids. The classifier achieved an accuracy of ~93% in 100 independent runs, with the data split into a training set (80% of the data) and a testing set (20%) (Figure 1D; Dataset S5). Two amino acid properties were most important in this classification (Figure 1E). The first is a low positive charge of amino acids in APRs compared to non-APRs. The second is a high amphiphilic propensity of these amino acids. The amphiphilic propensity measures the preference of amino acids to occur at protein-solvent interfaces, such as the end regions of transmembrane helices^32^. Amino acids like arginine and lysine have the highest preference for these environments, and the largest value of the amphiphilic index. In contrast, hydrophobic amino acids such as leucine, isoleucine, valine, and alanine prefer internal protein environments and have the lowest amphiphilic propensity^32^.

Because of the statistically significant difference between the amphiphilic nature of APRs and non-APRs, we argue that aggregator proteins resemble supramolecular amphiphiles. These molecular structures have distinct hydrophobic and hydrophilic parts^33^. In response to external stimuli such as temperature, pH, and ionic strength^33^, they can reversibly aggregate by non-covalent bonding forces, e.g., through electrostatic and hydrophobic interactions.

We next investigated the relevance of amphiphilic aggregation to stress-induced phase separation of our yeast aggregator proteins. An important feature of amphiphilic molecules is that they are more likely to occur in an aggregated phase if their hydrophobic chain is long^34^. We thus predicted that aggregator proteins with longer APRs will occur preferentially in the aggregated phase. To test this prediction, we asked whether aggregator proteins with long APRs preferentially occur in the pellet fraction of proteins extracted from heat-stressed yeast cells^35^. Specifically, we used for this purpose the log_2_ ratio of the protein’s abundance in the pellet fraction of yeast cells to that of supernatant as a measure of protein’s enrichment in the aggregated phase^35^. We found this enrichment data for 31 aggregated proteins in our dataset. Indeed, aggregator proteins with longer APRs preferentially accumulate in the aggregated pellet fraction (Figure 2A, R = 0.53, *p* = 0.0021; Spearman’s rank correlation). This association increases to R = 0.64 for aggregator proteins whose APRs have a low positive charge-to-aliphatic bias (p= 0.012; Spearman’s rank correlation). To see whether this observation extends to other stress-induced conditions, we also used the mass spectrometry dataset of nutrient-starved yeast cells reported by Narayanasamy et al^36^. We identified 56 aggregator proteins in this dataset and checked the association between APR lengths in these proteins and the Z-score of proteins’ enrichment in the pellet fraction. In nutrient-starved cells too, the aggregator proteins with longer APRs preferentially accumulate in the aggregated pellet fraction (Figures 2B, R = 0.47, *p* = 0.00014; Spearman’s rank correlation), and the association increased for the low positive charge-to-aliphatic bias (R = 0.53, *p* = 0.0022; Spearman’s rank correlation).

**Figure 2.**
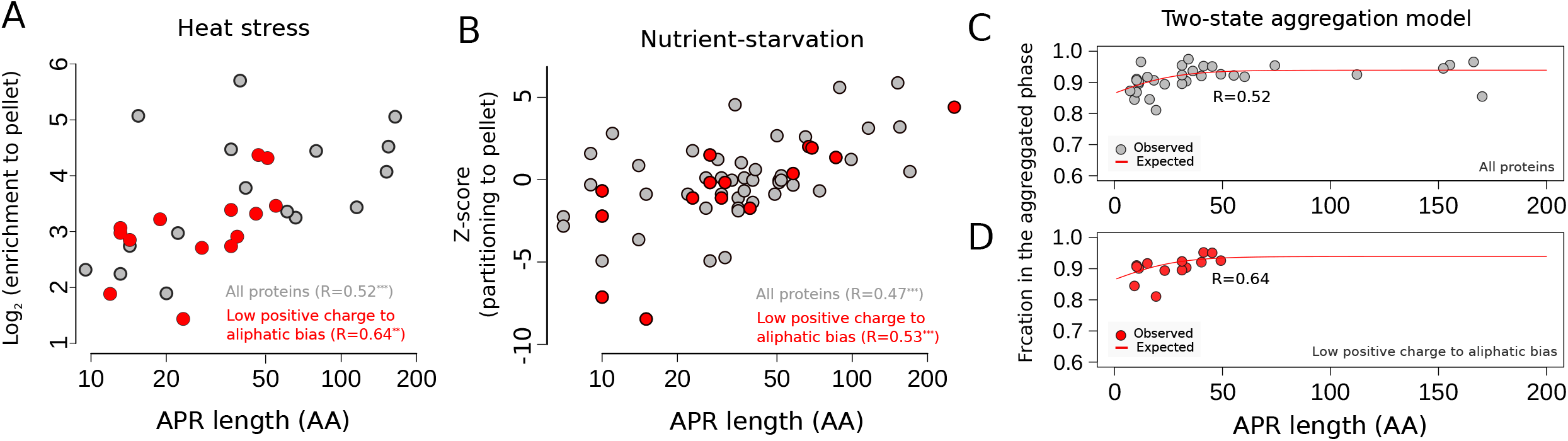
Aggregator proteins reversibly aggregate in a manner akin to amphiphilic molecules. A) The log2 ratio of the protein’s abundance in the pellet fraction to that of supernatant in heat-stressed yeast cells (30 out of 96 aggregator proteins^35^) versus their APR length. B) The aggregator proteins’ enrichment in the pellet fraction upon nutrient starvation (59 out of 96 proteins^35^) versus their APR length in log-scale. The red and the grey circles in both panels A and B represent all aggregator proteins, and those where the positive charge-to-aliphatic-bias in APRs was lower than its median value ~ 0.25. C) The fraction of aggregator proteins in the aggregated phase calculated from Equation 3 versus the length of APRs. D) The fraction of aggregator proteins with a low positive charge-to-aliphatic-bias in their APRs versus the length of APRs calculated from Equation 3. For both panels C and D, the fitted curve to the observed fraction of proteins in the aggregated phase using Equation 2 is shown in red. R values are the Spearman’s rank correlation between the expected (Equation 2) and observed (Equation 3) fractions of proteins in the aggregated phase.

We then investigated the relationship between APR length in aggregator proteins and their differential enrichment in the aggregated phase using equilibrium statistical thermodynamics. We considered a two-state model where proteins exist either as a soluble monomer or an insoluble aggregate. The fraction of proteins occurring in the aggregate at the critical concentration (cc) for aggregation 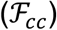 is expressed as:

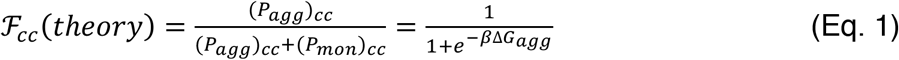

Here, *P_agg_* and *P_mon_* are the relative concentrations of proteins in the aggregated and the monomer phase, *β* = 1/*k_B_T* where *k_B_* is the Boltzmann’s constant, and Δ*G*_*agg*_ is the free energy change upon aggregation. Because the aggregation free energy increases in proportion to the length of the hydrophobic chain^37^, we rearranged Equation 1 to:

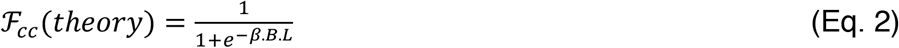

Where *B* is the proportionality constant between the aggregation free energy and the APR length. We then calculated the observed fraction of the aggregator proteins in the aggregated phase from their reported log_2_ enrichment ratio in the pellet fraction of heat-stressed yeast 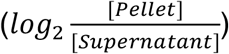 as:

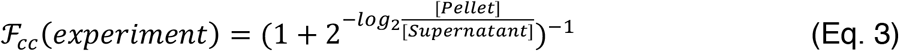

Equation 2 predicted the observed relationship between APR length and a protein’s fraction in the aggregated phase well (Figure 2C, R=0.53, p=0.0021; Spearman’s rank correlation, see supplementary information for the fitted constants). Importantly, the predictive power of Equation 2 was higher for proteins whose APRs had a low positive charge-to-aliphatic bias (Figure 2C; R=0.64, p=0.012; Spearman’s rank correlation).

In summary, our observations suggest that the aggregation-prone regions within the aggregator proteins can readily unfold in response to stressors, form hydrophobic long chains, and phase-separate in a manner akin to amphiphilic molecules. We also observed that aggregator proteins with longer APRs are more prone to aggregation. This relationship also suggests that aggregator proteins may leave the aggregated phase by masking their APRs from self-assembly or from interaction with other proteins. Molecular chaperones can facilitate this step: They have been co-purified with stress-induced aggregated proteins and can solubilize them^39^. Altogether, our study demonstrates that stress-induced phase separation is a sequence-encoded phenotype. It can be both explained and predicted by the self-assembly and aggregation of amphiphilic molecules.

## Methods

We identified aggregation-prone regions in the 96 aggregator proteins from the dataset of Cappelletti et. al^22^, and downloaded their sequences from the Uniprot database^40^ (Dataset S6). We chose 500 amino acid properties from the AAindex database to distinguish the physicochemical properties of APRs from non-APR segments in the aggregator protein^31^.

We used a binary classification to group regions within an aggregator protein into aggregation-prone regions (class A) and the remainder of the protein (class B). More specifically, we used a random forest classification, splitting aggregator proteins into a random subset comprising 80% of aggregator proteins for training and 20% for testing. To build features for the classification, we calculated the average value of 500 physicochemical properties for each sequence in the APR and non-APR dataset. This yielded two feature matrices for the APR and non-APR sequences. To apply random forest classification, we used the randomForest package in R^42^, and evaluated the best number of trees (*nTree*) and the number of variables randomly sampled at each split (*mtry*), in the random forest algorithm. To do so, we systematically varied the *nTree* and *mtry*, and calculated the accuracy of classification with 10-fold cross-validation and 3 repeats. We defined accuracy as the percentage of correctly identified classes of proteins (APRs and non-APRs) out of all instances. The combination of *nTree*=5000 trees and *mtry*=10 variables achieved the highest accuracy of ~ 90%. We then used these parameters to perform 100 random forest clusterings, in which we randomly assigned proteins to the training and the testing datasets. To quantify the accuracy of classification we counted the number of true positive and false positive predictions and calculated the area under the curve (AUC). These values are shown as a receiver operating characteristic curve (ROC) in Figure 2D. The most important physicochemical properties were the ones whose Gini index in the classification decreased the most compared to all other properties. This index is a widely-used measure of dispersion that reflects inequality in the values of a frequency distribution. Within the random forest framework, this index is calculated as the average probability that each of 500 physicochemical properties wrongly classifies APRs and non-APRs. We performed all models and statistical analyses using R. Scripts are available at: https://github.com/dasmeh/yeast_aggregators

